# StressNET: an adaptable deep-learning model for mechanical stress inference in tissues

**DOI:** 10.64898/2026.09.14.751456

**Authors:** Nicolás Aldecoa Rodrigo, Augusto Borges, Jerónimo R. Miranda-Rodriguez, Guillherme Ventura, Jakub Sedzinski, Hernán López-Schier, Osvaldo Chara

## Abstract

Mechanical interactions between cells are fundamental to tissue morphogenesis during development and regeneration. Computational methods that infer intercellular stresses from microscopy images of cell shapes offer a non-invasive alternative to experimental perturbation techniques, yet all existing approaches rely on explicit physical models. Here we present StressNET, a Graph Neural Network (GNN) that infers intercellular mechanical stresses directly from tissue geometry, without assuming any underlying physical model. We generated synthetic datasets to train and benchmark StressNET, and demonstrate that its predictions achieve state-of-the-art correlation with experimental stress proxies in zebrafish neuromasts and *Xenopus* embryos. Analysis of the network’s latent space reveals that StressNET learns global organizational principles of mechanical stress distribution, beyond local cell-cell interactions. StressNET is open-source and provides pre-trained models that can be fine-tuned on *in vivo* data, making it a broadly adaptable tool for studying tissue mechanics across biological systems.

## Introduction

Cellular interactions underpin biological diversity, with mechanical forces playing a key role in organizing cells into their final configurations. At the core of these interactions lie the mechanical properties of cells and their microenvironment, which control how forces are produced, sensed, and integrated^1–3^. Quantitative characterization of these properties is therefore essential for understanding morphogenesis.^4–7^. Mechanical parameters can be measured using various approaches, such as laser ablation ^8,9^, magnetic droplets ^10^, optical traps ^11^, or flipper probes ^12,13^. While these techniques have proved very instructive, they interfere with the biological tissue, are experimentally difficult, slow, or demand expensive equipment. Therefore, methods that estimate mechanical parameters solely from microscopy images, such as computational stress inference approaches, are non-destructive and provide an easier, faster, and cheaper solution^14–16^. Computational stress inference can be divided into two main categories: continuous methods, such as those that use a velocity field to estimate stress maps ^15^, and geometrical methods, which use the shape of cellular membranes in a tissue to estimate stress ^14,16^. Geometrical methods generally work by formulating the tissue mechanics as a series of linear equations, which are then solved simultaneously. These methods use approaches such as numerical inversion ^17–19^ , Bayesian solvers ^20^ and variational algorithms ^21^ to find solutions subject to certain physical rules. While most methods perform inference from a single image frame (static inference), other methods can estimate stress dynamically from image sequences ^16,19,22^.

Despite the wealth of biological information available through modern microscopy, a data-driven approach that leverages this data while assuming only that a tissue’s geometric arrangement encodes information about its internal stresses is lacking. Machine learning approaches provide a flexible alternative by learning complex, non-linear mappings directly from data ^23^. In particular, Graph Neural Networks (GNNs) are well-suited for this task, as they encode graph-structured data through iterative message passing between neighbouring elements, enabling the integration of local and global geometric features ^24–26^.

To address the lack of a data-driven approach to analyze tissue stresses from their geometric arrangement, here we present StressNET, a deep learning algorithm implemented as a GNN, which accurately predicts stresses in cellular membranes from microscopy images, as validated by *in silico* simulations, laser ablation experiments and myosin readouts. StressNET has four main components, which (1) extract edge features, (2) project them into a high-dimensional space, (3) aggregate the data across scales, and (4) obtain the predicted stresses through nonlinear embedding. We validated StressNET through simulations and *in vivo* experiments using Myosin II readouts as a proxy for stress, as well as laser ablation experiments in zebrafish neuromasts. We showed that StressNET accurately and precisely recapitulates the ground truth values. Importantly, we found that fine-tuning the network significantly improves accuracy with little additional training data, demonstrating that the StressNET algorithm can be adapted to new systems. Interestingly, by analysing the latent representation of the StressNET network, we observed that tissues cluster according to their pattern type, hinting at the ability of the network to learn broad organisation principles beyond local cell-cell interactions.

## Results

### StressNET predicts intercellular mechanical stresses from tissue geometry

StressNet is an automated graph neural network (GNN)-based pipeline that infers relative intercellular mechanical stresses directly from epithelial tissue images (Figure 1). Once images are acquired, the workflow consists of three stages: extraction of tissue geometry as a polygonal mesh, graph encoding of this geometry, and stress prediction using a GNN (Supplementary Results Section 1; Supplementary Figure 1). Segmented tissue images are skeletonised and converted into polygonal meshes, taking advantage of the ForSys software’s library ^19^, (Figure 1A–C), where each cell is represented as a polygon defined by ordered vertices. This representation is encoded as a graph in which nodes correspond to cellular membranes and edges relate connected cell junctions (Figure 1D). Stress inference is framed as a node-regression problem naturally suited to GNNs, as intercellular tensions depend primarily on local topology. Geometric information is encoded in adjacency, node-feature, and edge-feature tensors (Materials and Methods Section 1, Figure 1E–G), which serve as inputs for node-level prediction of relative stresses (Figure 1J, K).

**Figure 1.**
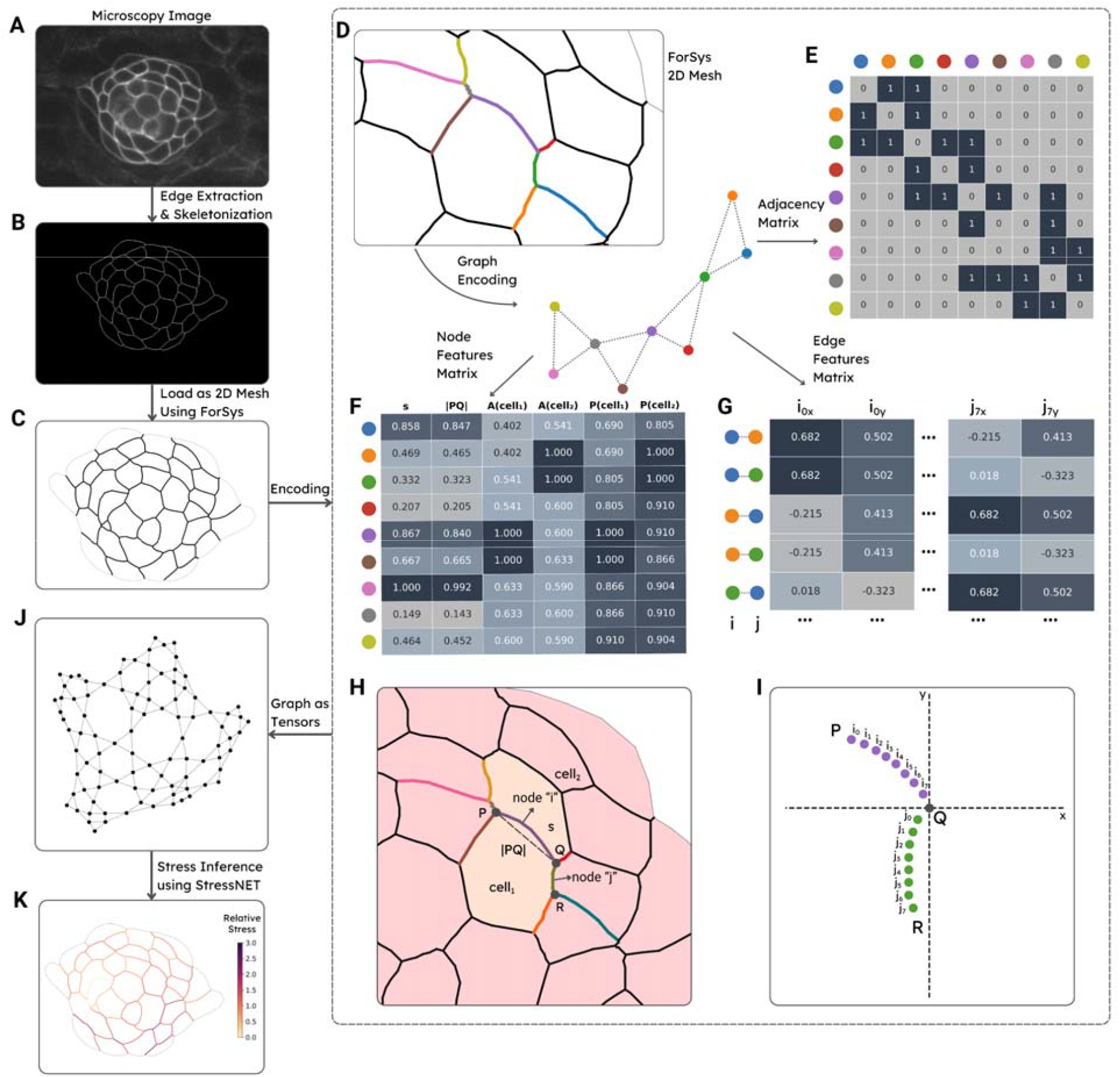
Workflow for inferring mechanical stress from microscopy images with StressNET. Our pipeline infers relative intercellular tensions from segmented microscopy images using a graph-based neural architecture. **(A)** A microscopy image of an epithelial tissue with membrane markers is used as input. **(B)** Tissue geometry is extracted using edge detection and skeletonisation. **(C)** The resulting contours are converted into a polygonal mesh using ForSys. **(D)** The mesh is represented as a graph, where nodes correspond to cellular membranes (polygon edges) and graph edges link a pair of nodes whose membranes share a junction. **(E)** The adjacency matrix encodes pairwise connectivity between interfaces. **(F)** The node-features matrix contains initial feature vectors describing each cell-cell interface. **(G)** The edge features matrix encodes the geometry between adjacent interface pairs. **(H)** Schematic of the features represented in the node-features matrix for node/interface “i” defined by the union of tricellular junctions P and Q. These features include the arc and chord lengths as well as the area and perimeters of its adjacent cells (Cell1 and cell2). **(I)** Illustration of the features contained at the edge feature matrix: the normalised relative coordinates of the interface PQ (i0:i7) and one of its converging interfaces, QR (j0:j7). **(J)** The three matrices together define a graph structure used as input to our model. **(K)** StressNET performs node-level regression to predict the relative stress magnitude at each interface.

To train and evaluate StressNet, we generated datasets comprising 14 stress-pattern families (100 samples each) using a 2D vertex model implemented in Surface Evolver through our seapipy pipeline ^19^. Five models with identical architectures and training procedures were trained independently using different random seeds, controlling weight initialisation, early stopping, and data augmentation. StressNet predictions were compared with our previous method, ForSys, in its static modality ^19^. StressNet accurately recovered ground-truth stress fields and, in some cases, outperformed ForSys (Supplementary Results Section 2; Supplementary Figure 2). Performance remained stable under noise added to stress patterns to mimic segmentation errors (Supplementary Results Section 3; Supplementary Figure 2). When evaluated on out-of-distribution data using single-prediction and augmented inference, StressNet generalised to topologies not seen during training (Supplementary Results Section 4; Supplementary Figure 3). Fine-tuning a subset of parameters on small datasets further improved performance on out-of-distribution simulations (Supplementary Results Section 5; Supplementary Figure 4), supporting application to experimental datasets where substantial domain shifts are expected.

### StressNet-predicted forces correlate with junction recoil velocity in zebrafish neuromasts

Following *in silico* validation, we applied StressNet to live imaging of zebrafish neuromasts, probing the junctional mechanical tension via targeted laser ablation. Using a membrane marker, we extracted cell geometries, performed single-junction ablations, and tracked the vertices delimiting each junction. Recoil velocities were obtained from linear fits to post-ablation vertex separation and, under an overdamped regime, report the pre-ablation tensile load, providing an experimental benchmark for predicted relative stresses ^27^.

We quantified correlations between predicted stresses and recoil velocities across 15 neuromasts whose inference showed a condition number of less than 500 (For details, see Materials and Methods Section 2; and Supplementary Figure 5) (Figure 2A–C). Progressive exclusion of recoil estimates with poorly conditioned linear fits led to a monotonic increase in correlation for StressNet predictions, highlighting sensitivity to numerical stability (Materials and Methods Section 2; Supplementary Figure 5). The final dataset showed a moderate correlation between ForSys, used here as an independent reference stress-inference method, and recoil velocity (Pearson r = 0.548, p = 0.035; Figure 2D), whereas the five StressNet base models showed substantially stronger correlations (mean Pearson r = 0.813 ± 0.057; all p < 0.003; Figure 2E), consistently outperforming this baseline (Figure 2F). These results demonstrate that StressNet captures stress differences reflected in *in vivo* recoil responses despite being trained exclusively on synthetic data, motivating its fine-tuning to further improve performance across untested epithelia.

**Figure 2.**
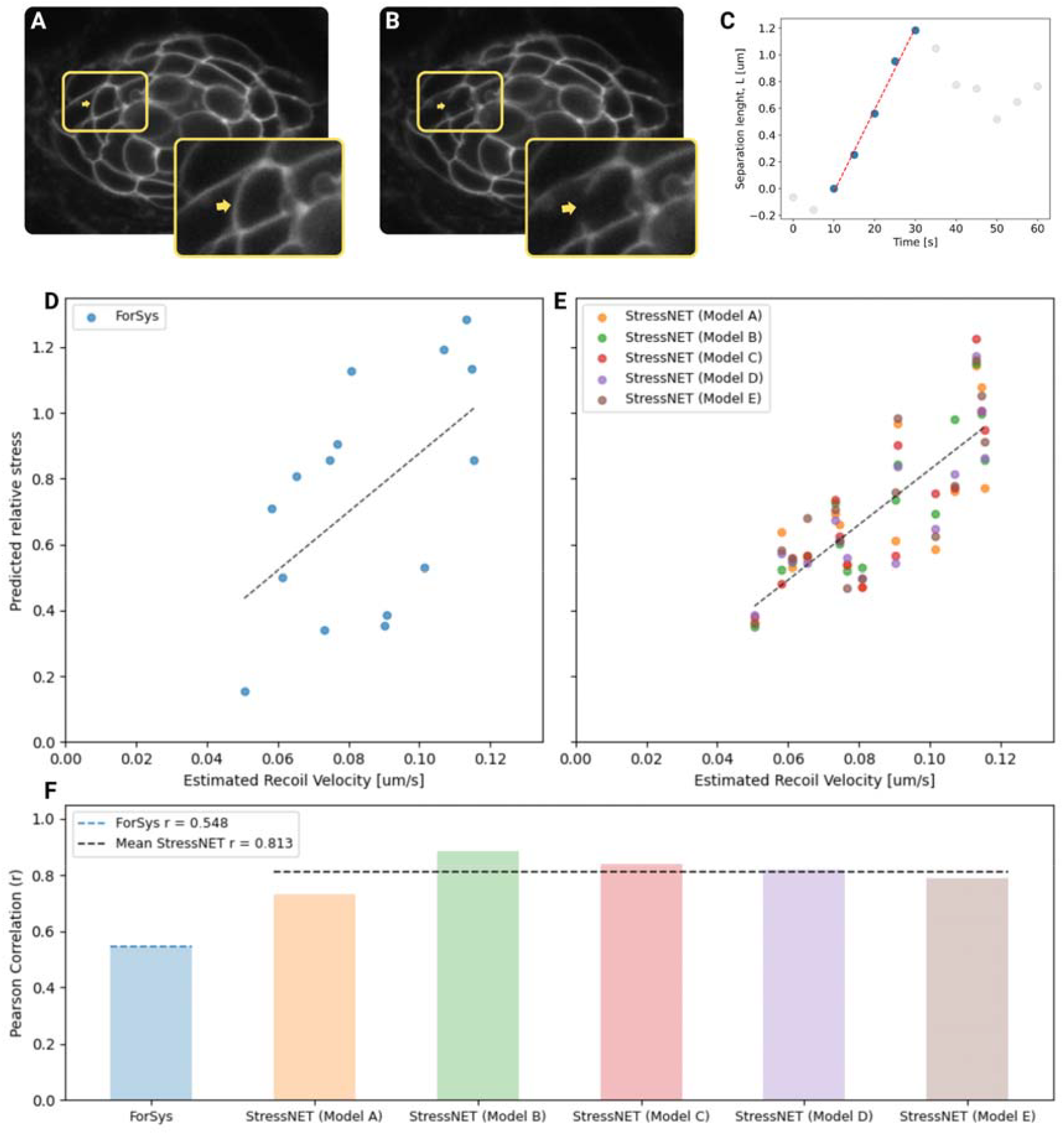
Predicted tensions correlate with recoil-velocity measurements from single-junction laser ablations. **(A)** Microscopic image of a neuromast prior to laser ablation. The inset provides a magnified view of the targeted cell-cell interface, indicated by the yellow arrow. **(B)** Corresponding image immediately after ablation, with the inset highlighting the severed junction. **(C)** Example displacement-time trace for the two recoiling vertices, along with the linear fit used to estimate the initial recoil velocity. **(D)** Predicted relative stresses from the ForSys model plotted against estimated recoil velocities for a curated set of 15 laser ablations. The dashed line indicates the linear fit summarising the overall trend. **(E)** Same analysis as in (D), but showing predictions from five independently trained StressNET base models. The dashed line represents the linear fit computed across all StressNET predictions combined. **(F)** Correlation analysis comparing predicted relative stress with recoil velocity for ForSys and each StressNET model. Bars indicate the Pearson correlation for each model between the predicted relative stress at the ablated edge and the corresponding recoil velocity. The blue dashed line denotes the correlation obtained with ForSys, while the black one shows the average correlation of the five StressNET models.

### StressNet inferences correlate with Myosin II localisation at epithelial junctions in Xenopus embryos and can be boosted through fine tuning

We next applied StressNet to an independent *in vivo* system: mucociliary epithelia of Xenopus embryos expressing the SF9-3xGFP myosin II sensor, which reports non-muscle myosin IIA enrichment at epithelial junctions ^28^. This system provides a quantitative readout of myosin-driven contractility that can be directly compared with inferred stress values (Materials and Methods Section 3, Figure 3A). Across 126 images, models trained exclusively on synthetic data showed significantly higher correlation with myosin intensities than the ForSys baseline (p=1.6×10^−11^); median Pearson correlation values were 0.39 for StressNET and 0.18 for ForSys (Table 1, Figure 3B).

**Figure 3.**
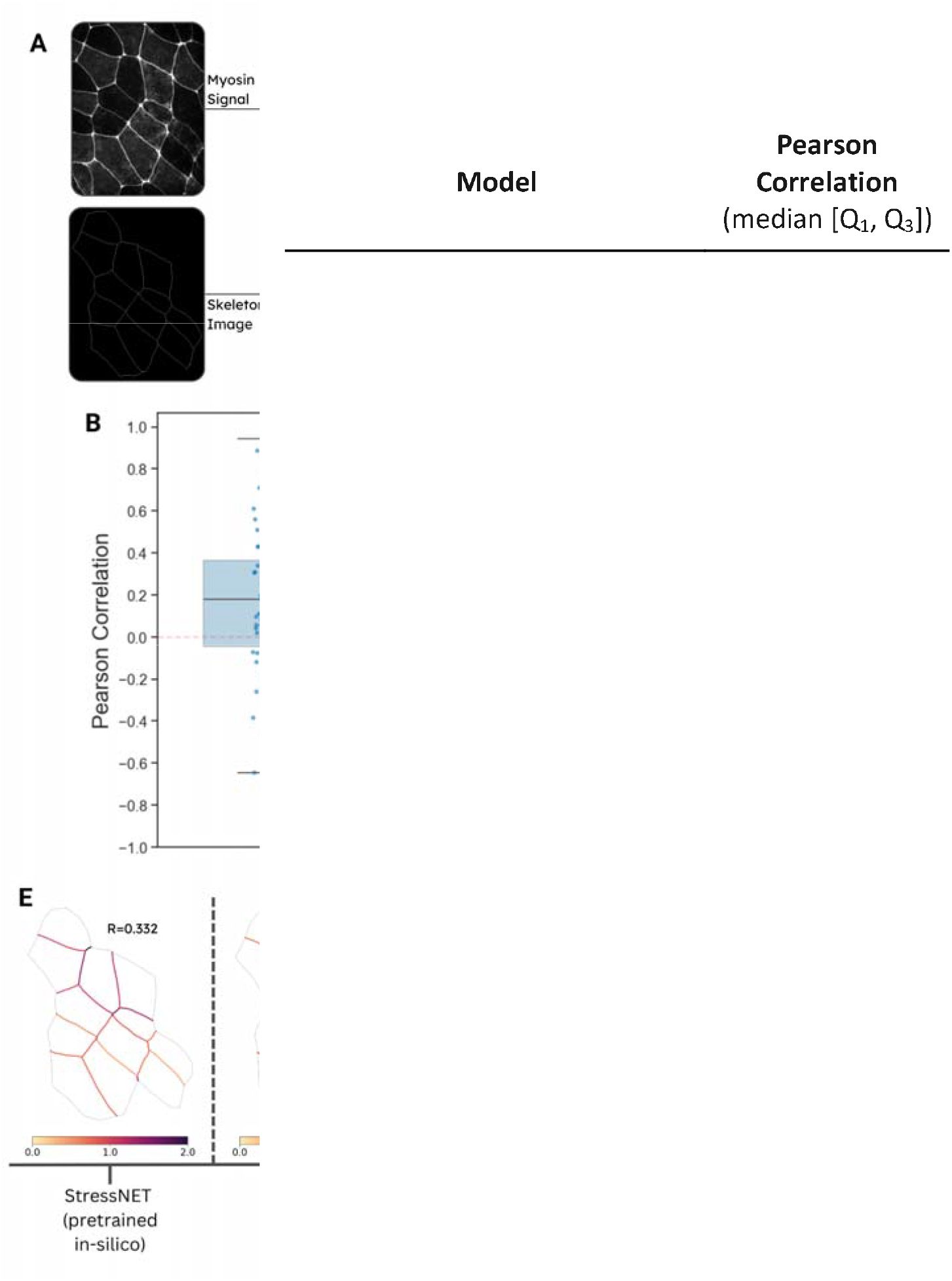
StressNET predicts *in vivo* Myosin II intensity at cell junctions on unseen data and is significantly improved by minimal fine-tuning. **(A)** Experimental validation workflow: microscopy images are skeletonised and converted into a 2D lattice. Ground-truth cell-cell interface tensions are assigned from normalised Myosin II signal intensity. **(B)** Pearson correlation between predicted relative tensions and normalised myosin intensity across test samples, showing that pretrained StressNET models exhibit significantly stronger alignment than ForSys. Boxplot whiskers extend to the most extreme data points within 1.5× the interquartile range. **(C)** Fine-tuning configuration: only approximately 2.28% of parameters were updated, while all other weights in pre-trained StressNET models remained fixed. **(D)** The mean Pearson correlation between predicted tensions and experimental myosin II intensities exhibits a logarithmic dependence on the number of fine-tuning samples, with shaded regions denoting confidence intervals derived from bootstrap resampling. The inset presents the linear regression on the mean correlations computed in the log-transformed sample space. **(E)** Visualisation of tension maps from an example in the evaluation subset, demonstrating closer alignment with normalised myosin II signal intensity as the amount of fine-tuning data increases.

**Table 1:** Pearson correlation between predicted tensions and normalised myosin II fluorescence intensity in Xenopus mucociliary epithelium images, comparing StressNET and ForSys. Correlations were calculated on the full dataset (126 images) for pretrained models and the ForSys static method, and on a held-out subset (38 images) for pretrained and fine-tuned StressNET models as well as ForSys. Fine-tuning updated a small fraction (approximately 2.28%) of StressNET parameters with increasing sample sizes, resulting in significant performance gains. Fine-tuned StressNET with augmented inference achieved moderate-to-high correlations, substantially outperforming existing approaches.

|  |  |
| --- | --- |
| ForSys<br>static | 0.18 [-0.04, 0.37]<br>(n=126) |
| **StressNET**<br>**pre-trained *in silico*** | **0.39 [0.23, 0.54]**<br>(n=126) |
| ForSys<br>static | 0.12 [-0.18, 0.29]<br>(n=38) |
| StressNET<br>pre-trained *in silico* | 0.37 [0.11, 0.50]<br>(n=38) |
| StressNET<br>fine-tuned *in vivo* (88 samples)<br>single prediction | 0.58 [0.45, 0.70]<br>(n=38) |
| **StressNET**<br>**fine-tuned *in vivo* (88 samples)**<br>**augmented prediction** | **0.59 [0.47, 0.73]**<br>(n=38) |

To assess the benefit of fine-tuning, models were trained on progressively larger subsets of the dataset and evaluated on an independent test set of 38 images (Figure 3C). Performance increased consistently with training set size, with the average Pearson correlation exhibiting a logarithmic dependence (R^2^ > 0.97, Figure 3D-E), consistent with trends observed *in silico* (Supplementary Figure 4B). Fine-tuning on 88 samples yielded a median Pearson correlation of 0.59 on the test set (Table 1). Even minimal fine-tuning substantially improved performance: using only 11 randomly selected samples increased the mean Pearson correlation from 0.29 to 0.44 (single prediction) and 0.48 (augmented), representing an improvement of ∼50%. These results demonstrate that StressNet predictions can be efficiently adapted to new experimental datasets with minimal additional training data, validating its generalizability across diverse biological systems.

### StressNet’s latent space encodes global cues, such as the spatial distributions of stresses

StressNet predicts global tissue stresses by aggregating local node features into a context-enriched latent representation. To test whether this representation captures broader organisational principles, we analysed Global Attention Sum Pooling embeddings. These embeddings provide a global latent representation of the tissue, obtained by attention-weighted aggregation of node features. The embeddings were generated from newly simulated data from the 14 in-distribution spatial pattern families, excluding training samples (Materials and Methods Section 4). Linear probes showed that graph size (number of cell-cell interfaces) is largely recoverable from the embeddings (R^2^ = 0.943 ± 0.009 vs –0.215 ± 0.013 for permuted controls). Pattern family could be predicted with an accuracy of 0.856 ± 0.021, compared to 0.0702 ± 0.001 for permuted labels, the latter being consistent with random guessing across 14 classes (Supplementary Figure 6).

PCA of z-score standardized embeddings showed that PC1 separates spatially independent (“non-local”) from spatially continuous (“local”) stress patterns, while PC2 correlates with graph size (point-biserial correlation = 0.810 ± 0.090; Spearman ρ = 0.750 ± 0.066; all p < 1×10^−16^), increasing to 0.849 ± 0.039 when excluding non-local patterns (Figure 4A(i), B(i)). When projecting out-of-distribution simulations and in vivo Xenopus and neuromast tissues onto the principal components defined by the in-distribution patterns, PC1 remained stable for the synthetic families, which all occupy the spatially coherent (“local”) regime. In vivo samples showed consistent alignment with this axis, with neuromast tissues clustering within the local regime and a subset of small Xenopus systems shifted towards more non-local configurations (Figure 4A(ii–iii)). PC2 correlated with graph size (Spearman ρ = 0.789 ± 0.078; all *P* < 1 × 10^−16^), consistent with observations on in-distribution patterns (Figure 4B(ii–iii)).

**Figure 4.**
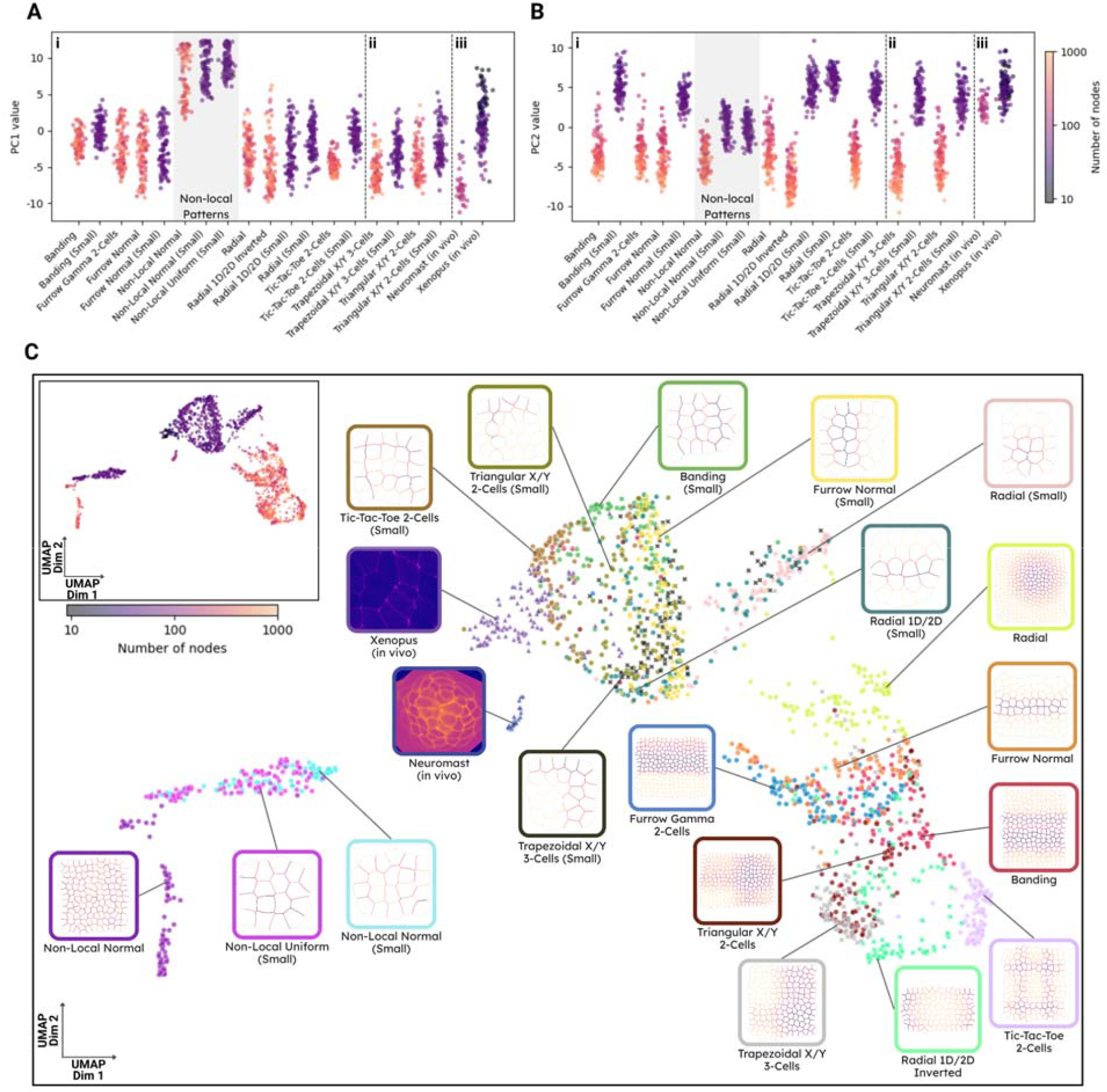
StressNET’s latent space encodes global cues and reveals the underlying spatial patterns of stresses. **(A–B)** Principal component analysis (PCA) of Global Attention Sum Pooling embeddings from one representative StressNET model (Model C) reveals two dominant, interpretable axes of variation. Each point represents a tissue configuration. Panels are divided into three sections *(i-iii)*, separated by dashed lines, corresponding to *(i)* synthetic pattern families used during training, *(ii)* held-out synthetic families, and *(iii)* in vivo samples; with PCA fitted on training-distribution embeddings and applied to all samples. **(A)** The first principal component (PC1) separates non-local from spatially coherent pattern families. This organization is consistent across all groups, with in vivo samples largely aligning with spatially coherent (local) patterns, except for a subset of small Xenopus tissues (<10 nodes) that are shifted toward non-local configurations along this axis. **(B)** The second component (PC2) correlates strongly with the number of nodes in each graph, indicating that StressNET’s latent representations capture system size. Points are coloured by the number of nodes (on a logarithmic scale). **(C)** UMAP projection of the same embeddings across all samples, colored by pattern family and annotated with representative examples showing cell-cell interfaces colored by ground-truth tension for synthetic samples and microscopy images for experimental data; marker shape indicates data source, with circles for training families, crosses for held-out families, and triangles for in vivo samples. The inset (top left) shows the same projection colored by the number of nodes (logarithmic scale). The embedding space reveals a hierarchical organization, with a clear separation between local and non-local patterns, further structured by system size and clustering across pattern families and in vivo data.

UMAP visualisation revealed a distinct cluster of non-local spatial patterns and a continuous organisation of samples structured by graph size, with consistent separation across pattern families. Out-of-distribution simulations and in vivo samples occupied coherent regions in the two-dimensional projection, with neuromast tissues occupying a compact region of the embedding space and Xenopus samples forming a less well-defined cluster, in the periphery of synthetic local patterns clusters (Figure 4C). Together, these results indicate that StressNet embeddings encode global structural and contextual properties of the tissue despite the model being trained only for node-level relative stress regression, suggesting that such contextual mechanical information is essential for prediction performance. The emergent structure observed in the learned representation can be interpreted as a *forcespace*.

## Discussion

Mechanical interactions between cells are at the heart of how tissue shape emerges ^2,3,29,30^. Mechanical stress inference provides an inexpensive and straightforward computational method for determining the mechanical state of a tissue from the geometry of cellular plasma membranes ^14,16^. StressNET is an automated graph neural network (GNN)-based pipeline that enables non-invasive estimation of the mechanical state of tissues where cell boundaries are visible, using only segmented microscopy images as an input. Currently, all computational stress inference methods encode the physical constraints on the system as a system of equations, which are then simultaneously solved to infer the stresses ^14,16^. StressNET is the first deep learning-based approach which, instead of creating a system of equations to represent the morphologies observed in microscopy images, it builds a Graph Neural Network (GNN) that integrates local and global tissue information.

One of the main limitations of computational stress inference is its dependence on the segmentation quality of the raw microscopy images ^14,16^. We have shown that perturbations in membrane shape, induced by jittering of the vertices, degrade accuracy less when comparing StressNET with other methods. As a deep learning algorithm, StressNET benefits from data augmentation, which increases its accuracy at the cost of further training.

One key advantage of deep learning models is their ability to adapt learned representations to the specifics of a given task through fine-tuning, thereby refining previously learned features for new domains ^31–33^. In the mucociliary epithelium of Xenopus, we used Myosin II as a proxy for cortical stress and found that StressNET’s predictions correlated with Myosin II intensity at the membranes. After fine-tuning, StressNET’s predictions showed a logarithmic improvement, increasing from R=0.3 to R=0.6. This suggests that StressNET can be adapted to different experimental systems through fine-tuning, by using a small number of microscopy images.

We then turned to the latent representation generated by our network. We studied the global attention pooling vector using two dimensionality reduction approaches: PCA (Principal Component Analysis) and UMAP ^34^ (Uniform Manifold Approximation and Projection). Both approaches showcased StressNET’s ability to discriminate local and non-local patterns, suggesting that the network uses global tissue information to generate local stress predictions and learns general mechanical rules. GNNs have already been applied to cultured MCF-10A cells ^35^ and the Drosophila ventral furrow ^36^ to predict the relationship between cell geometry and dynamics. GNNs have been shown to predict different aspects of morphogenesis during Drosophila gastrulation at the bulk, edge, and node levels ^37^. Moreover, GNNs were used in the basal layer of the epidermis to predict cellular fate by training on local cell features such as cell contacts or gene expression ^38^. These works and ours suggest that GNNs could prove useful in studying and predicting tissue and cellular events by learning and generalising developmental and morphogenetic rules.

In the future, adapting the software to three dimensions would be critical to enable volumetric data analysis of more complex systems. This could be done by extending the feature representation to three-dimensional lattices, allowing StressNET to leverage the same architecture while capturing additional geometric relationships. Similarly, StressNET could be adapted to time-dependent stress inference ^16,19^ by incorporating temporal information across system geometry snapshots, enabling the model to predict dynamic stress patterns. The *forcespace* generated through StressNET could also be used in combination with other observables of the system, such as shape (e.g., morphospace) ^39^ or omics data ^40^, to understand how different levels of information, transcriptional and mechanical, are integrated to generate robust tissue shape.

The ability to infer mechanical stresses non-invasively has opened new avenues for studying tissue mechanics. StressNET’s capacity to learn global tissue properties and translate them into local stresses suggests that shared geometric principles underlie intercellular stress organization across diverse contexts. As imaging technologies continue to develop, tools like StressNET will be essential for translating the wealth of microscopy data into quantitative mechanical observations, bridging the gap between tissue geometry and the mechanical programmes that drive morphogenesis.

## Materials and Methods

### 1. Overview of StressNET’s model and inference workflow

#### Data processing steps

StressNET requires a segmented skeletonised representation of the tissue of interest. Given the availability of robust tools for edge detection and skeletonisation ^41^ , extracting clean representations of cell boundaries has become a well-established preprocessing step ^17,19,42^.

From the segmentation, the geometrical properties of the system are encoded in the adjacency matrix, the node-feature matrix, and the edge-feature matrix, as noted in the Results section.

The adjacency matrix *(Figure 1E)* connects nodes in the graph bidirectionally. Connected nodes correspond to cell-cell interfaces that share a vertex (junction point).

The Node Features matrix (Figure 1F) has size N×6, where N represents the number of cell-cell interfaces in the system after excluding external and disconnected ones. Each node is represented by a six-dimensional feature vector encoding engineered attributes (Figure 1H). The first feature is the arc length, calculated as the sum of Euclidean distances between consecutive vertices outlining the interface; the second is the chord length, computed as the euclidean distance between the two endpoints of the interface; the third and fourth features correspond to the areas of the two cells sharing the interface; and the fifth and sixth features represent the perimeters of these two cells. For interfaces that do not have two complete cells defined within the lattice, the available cell’s area and perimeter values are duplicated to fill the corresponding pair of features, ensuring a consistent six-dimensional representation. The arc length and chord length features in this matrix are divided by the maximum arc length across all interfaces, preserving their relative scale while ensuring values remain below 1. Cell areas and perimeters are each divided by their respective maxima, ensuring all feature values remain below 1.

The Edge Features matrix (Figure 1G) encodes the geometry of pairs of cell-cell interfaces that converge at junction points. For each edge connecting nodes “i” and “j,” the matrix stores the 2D coordinates (x, y) of points along both interfaces (flattened into a one-dimensional vector), with the coordinates of node “i” listed first, followed by those of node “j” (Figure 1I). These coordinates are normalised by subtracting the junction point position and dividing by the maximum arc length from the Node Features matrix, to ensure that all values lie within the range [-1, 1], with the junction point fixed at (0, 0). The constant junction point coordinates are excluded from the matrix to avoid introducing spurious parameters, resulting in a tensor of shape E × 4(P−1) (in our experiments, P = 9), where E represents the number of edges (i.e., the number of 1s in the adjacency matrix) and P is the number of points defining each interface. When the input data contain a different number of points, vertices are resampled to a fixed length using quadratic spline interpolation, ensuring a consistent tensor shape across samples. To maintain consistency with the adjacency matrix and the Edge Features tensor, the latter is sorted in row-major order, as required by the Spektral library ^43^ used to implement the GNN layers in our model.

The output is a vector containing the predicted relative stress magnitudes at each cell-cell interface. To facilitate comparison with other stress inference approaches that estimate edge tensions under a mean constraint ^17,19,20,42^, the predicted values are normalised such that the average stress value is equal to one.

#### Neural network architecture

StressNET is composed of four main components: an edge-feature extractor that encodes geometric information from edge-coordinate sequences; a deep graph neural network backbone built from StressConv layers that propagates information across the graph; a node-attention global pooling module that computes a graph-level descriptor; and a feedforward regression head that predicts node-level relative tensions. A complete layer-by-layer description of the architecture, including inputs, output shapes, trainable parameters, and their contribution to the total parameter count, is provided in *Method’sTable 1*.

The edge-feature extractor (Supplementary Figure 1A) is a feedforward network composed of two dense layers, also known as fully connected (FC) layers ^44^. Each layer is followed by layer normalisation ^45^, which standardises the activations across each input to stabilise and accelerate training. A non-linear activation function, specifically the Exponential Linear Unit (ELU) ^46^, is applied immediately before the second FC layer. The first layer projects the input into a higher-dimensional space, allowing the network to model non-linear relationships between features. The second layer reduces the representation to a smaller vector, acting as a bottleneck that forces the network to learn a more compact representation. This reduction decreases the computational cost of subsequent components compared to maintaining a high-dimensional representation and may act as a regularizer, constraining the model to produce more robust features ^47,48^.

The input to the network is the raw sequences of 2D coordinates encoded for each graph edge “ij”, flattened into vectors of length 2P. In our final model, this component outputs an E × 128 matrix of extracted edge features, which are directly fed into each StressCONV layer without further modification. With a latent vector dimensionality of 512, this component accounts for approximately 11% of the total model parameters.

The backbone of the StressNET model is a 12-layer graph convolutional network ^25^ organised into four distinct blocks (Supplementary Figure 1B), each containing three StressConv layers of the same width, with the width increasing progressively from one block to the next. These blocks are connected sequentially. The first layer of each block, a “transition” layer, increases the dimensionality of the node representations, while the subsequent “intermediate” layers maintain this dimensionality constant throughout the block.

The StressConv architecture, which we developed, is inspired by the CrystalConv layer available in the Spektral library and can be seen as an extension of the formulation by Xie and Grossman ^49^.

StressConv updates each graph node based on its local neighbourhood. For a given node “i”, the following operations are applied:

1. The feature vectors of the node *i*, each neighbour *j*, and the corresponding edge e_ji_ are concatenated into a single vector to form pairwise representations, which are then projected into a higher-dimensional space by multiplying them with a weight matrix W^(a)^. Afterwards, layer normalisation is applied to stabilise training, followed by the Exponential Linear Unit (ELU) activation function (Eq. S1). In our experiments, this higher-dimensional representation is set to be twice the size of the layer output.

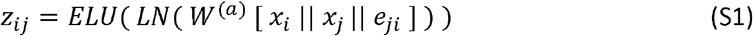

Where ELU is the Exponential linear unit, LN is the layer normalisation (centre and scaled), W^(a)^ is the weight matrix and , , and are the feature vectors for nodes *i* and *j* and the edge that connects from *j* to *i*. The “||” operator is vector concatenation.
2. The output from the previous step is passed through two fully-connected (FC) sublayers, each with its own set of weights and biases. Sublayer “b” performs a linear down-projection of the latent representation, following the same approach used in the Edge Feature Extractor MLP. Sublayer “g” computes a tensor of gating coefficients with the same shape. The sigmoid activation function is then applied to constrain these coefficients between 0 and 1. The output of the “g” sublayer is then element-wise multiplied by the output of the “b” sublayer (Eq. S2). This modulation enables the network to selectively emphasise or attenuate specific features in the joint representation of nodes and their neighbours before aggregation.
3. For each node, the resulting feature vectors from the node and its neighbours are aggregated into a single representation by summing them element-wise (Eq. S2).

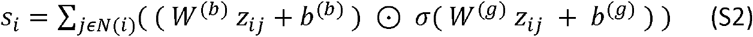

Where *N(i)* are the neighbours of node *i*, z_*ij*_ is the output from the previous step for a given self-neighbour pair, *b* are the bias vectors, ⊙ is the element-wise multiplication, and σ-is the sigmoid activation function
4. Finally, an “add and norm” ^50^ operation creates a residual connection with the output of the previous layer and standardises the resulting tensor by applying layer normalisation (Eq. S3). In Transition StressConv Layers, where the input and output dimensions differ, the input is linearly projected by multiplying it by an additional learned weight matrix W^(r)^ before addition to match the tensor shapes (Eq. S4).

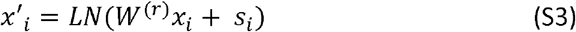

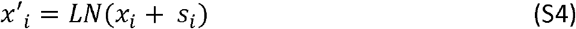

Where are the updated feature vectors for node *i* after normalisation.

The GNN integrates both edge and node features to progressively enrich and refine their representations, ultimately producing a 96-dimensional vector for each node to which the ELU activation function is applied to obtain the final node feature vector. Each graph convolution layer directly relates a node’s features to those of its neighbours. However, as additional layers are applied, the receptive field expands, indirectly incorporating information from the neighbours of neighbours. After 12 layers, the effective receptive field is large enough to enable message passing across substantial sections of the graph.

The output widths for the StressConv layers in the four blocks are set to 32, 48, 64, and 96, respectively. Under these settings, the Deep GNN component accounts for almost 80% of the model’s total parameters.

Node features are aggregated into a global descriptor (Supplementary Figure 1C) using a node-attention global pooling layer, which learns attention coefficients to calculate weights for the nodes and sum their features. We utilised the ‘GlobalAttnSumPool’ implementation from the Spektral package for this purpose. This layer learns a vector of weights that matches the length of each node’s feature vector, which is 96 in our case. It computes the dot product between the learned weights and the node-features matrix and applies the softmax function to obtain attention coefficients for the nodes that sum to 1. These coefficients are then used to pool node features by performing a weighted sum of all vectors. The pooled feature vector is then concatenated with each node feature vector, resulting in a final representation that enables the network to relate each node’s features to a graph-level descriptor before performing the regression of relative tensions. Inspired by the approach used in PointNet ^51^ the addition of this global descriptor reduces the need for the GNN to condense extensive contextual information into single node representations, allowing for a more effective distribution of both local and global features.

The final step consists of a non-linear regression performed by a feed-forward network with three FC layers (Supplementary Figure 1D), sharing weights across nodes. This is followed by the MeanNorm layer, which normalises the output vector by dividing it by its mean. The first layer projects features to 256 dimensions and applies layer normalisation along with ELU non-linearity. The second layer reduces the dimensionality to 64, with another layer of normalisation and ELU activation applied. Finally, a linear FC layer with an output size of one produces a scalar value, which is interpreted as the node’s relative tension. This component accounts for approximately 9% of the model’s total parameters.

**Methods Table 1.** Layer-by-layer summary of the StressNET graph neural network. The table lists each layer, its inputs, output shape, number of trainable parameters, and the percentage contribution to the total parameter count. Inputs include node features (node_feats, ⍰×6), the adjacency matrix (adj_mat, ⍰×⍰), and edge features (edge_feats, ⍰×32), where ⍰ and ⍰ denote the number of nodes and edges. Parameter counts correspond to trainable weights, and percentages indicate each layer’s share of the total parameter count.

| Layer | Inputs | Output shape | # parameters | % parameters |
| --- | --- | --- | --- | --- |
| node_feats | - | (N, 6) | 0 | 0 |
| adj_mat | - | (N, N) | 0 | 0 |
| edge_feats | - | (E, 32) | 0 | 0 |
| edge_feats_fc_1 | edge_feats | (E, 512) | 16384 | 2.25 |
| edge_feats_ln_1 | edge_feats_fc_1 | (E, 512) | 1024 | 0.14 |
| edge_feats_elu_1 | edge_feats_ln_1 | (E, 512) | 0 | 0 |
| edge_feats_fc_2 | edge_feats_elu_1 | (E, 128) | 65536 | 9.01 |
| edge_feats_ln_2 | edge_feats_fc_2 | (E, 128) | 256 | 0.04 |
| graph_conv_1.1 | node_feats, adj_mat, edge_feats_ln_2 | (N, 32) | 13504 | 1.86 |
| graph_conv_1.2 | graph_conv_1.1, adj_mat, edge_feats_ln_2 | (N, 32) | 16640 | 2.29 |
| graph_conv_1.3 | graph_conv_1.2, adj_mat, edge_feats_ln_2 | (N, 32) | 16640 | 2.29 |
| graph_conv_2.1 | graph_conv_1.3, adj_mat, edge_feats_ln_2 | (N, 48) | 29568 | 4.07 |
| graph_conv_2.2 | graph_conv_2.1, adj_mat, edge_feats_ln_2 | (N, 48) | 31104 | 4.28 |
| graph_conv_2.3 | graph_conv_2.2, adj_mat, edge_feats_ln_2 | (N, 48) | 31104 | 4.28 |
| graph_conv_3.1 | graph_conv_2.3, adj_mat, edge_feats_ln_2 | (N, 64) | 48640 | 6.69 |
| graph_conv_3.2 | graph_conv_3.1, adj_mat, edge_feats_ln_2 | (N, 64) | 49664 | 6.83 |
| graph_conv_3.3 | graph_conv_3.2, adj_mat, edge_feats_ln_2 | (N, 64) | 49664 | 6.83 |
| graph_conv_4.1 | graph_conv_3.3, adj_mat, edge_feats_ln_2 | (N, 96) | 92928 | 12.78 |
| graph_conv_4.2 | graph_conv_4.1, adj_mat, edge_feats_ln_2 | (N, 96) | 99072 | 13.62 |
| graph_conv_4.3 | graph_conv_4.2, adj_mat, edge_feats_ln_2 | (N, 96) | 99072 | 13.62 |
| graph_conv_out_elu | graph_conv_4.3 | (N, 96) | 0 | 0 |
| global_pool | graph_conv_out_elu | (1, 96) | 96 | 0.01 |
| concat_global_feats | graph_conv_out_elu, global_pool | (N, 192) | 0 | 0 |
| reg_head_fc_1 | concat_global_feats | (N, 256) | 49152 | 6.76 |
| reg_head_ln_1 | reg_head_fc_1 | (N, 256) | 512 | 0.07 |
| reg_head_elu_1 | reg_head_ln_1 | (N, 256) | 0 | 0 |
| reg_head_fc_2 | reg_head_elu_1 | (N, 64) | 16384 | 2.25 |
| reg_head_ln_2 | reg_head_fc_2 | (N, 64) | 128 | 0.02 |
| reg_head_elu_2 | reg_head_ln_2 | (N, 64) | 0 | 0 |
| reg_head_out | reg_head_elu_2 | (N, 1) | 65 | 0.01 |
| mean_norm | reg_head_out | (None, 1) | 0 | 0 |

### 2. Recoil velocity estimation and dataset curation in zebrafish neuromasts

#### Laser ablation experiments

Laser ablation experiment data were generated in Borges et al. and reanalysed for this paper ^19^. Zebrafish larvae carrying Tg[−8.0cldnb:Lyn-EGFP] ^52^ were anaesthetised in MS222 and mounted in low– melting-point agarose. Neuromasts located in positions L2 or L3 were imaged using a custom-built Zeiss spinning-disk confocal microscope with a 63× water-immersion objective. For each ablation experiment, a single junction was severed using a pulsed 355-nm iLasPulse laser (400 ps), with three pulses delivered between the third and fourth frame. Time-lapse recordings were acquired at 1 frame per second, and the two tricellular vertices delimiting the ablated junction were manually tracked in FIJI.

#### Recoil velocity estimation and dataset curation

The Euclidean distance between the tracked vertices was computed over time, and recoil velocities were obtained by ordinary least-squares fitting of the post-ablation separation trajectory. For each fit, we extracted the slope, its standard error, the associated p-value, and the condition number of the covariance matrix returned by the optimiser. These quantities were used downstream to curate the dataset and define numerically stable subsets for quantitative comparison with model predictions.

Cellular geometries for tension inference (junction identities and lengths) were obtained by segmenting membrane-marker images with ilastik ^53^ (autocontext model followed by the multicut workflow), importing the resulting skeletonisations into TissueAnalyzer ^54^ for manual correction and edge tracking, and exporting the final cell–junction graphs. These graphs were used as input to both StressNET and the ForSys baseline, enabling direct comparison of predicted tensions with the experimentally measured recoil velocities.

We began with 37 laser-ablation experiments. Recoil velocities were estimated by linear regression of the post-ablation displacement. Experiments in which the fitted slope was not statistically significant (p > 0.05) were excluded, as these cases do not provide a reliable measure of mechanical recoil. This removed two samples.

We next applied an outlier-detection procedure to ensure that the tension-recoil relationship was not driven by individual inconsistent measurements. Outliers were identified independently for the ForSys baseline and for each of the five StressNET models using a two-dimensional Mahalanobis distance criterion on the paired tension–recoil values, with a χ^2^ threshold corresponding to p = 0.05. Only samples flagged as outliers in all six analyses were removed; this step eliminated two additional experiments, leaving 33 for subsequent analysis.

Because the reliability of the recoil-velocity estimate depends on the numerical stability of the linear fit, we generated progressively more stringent subsets using the condition number of the resulting covariance matrices as a criterion. We computed correlations using the full curated set of 33 experiments, and then generated progressively more stringent subsets by restricting the condition number of the covariance matrix to values below 2000, 1000, and 500, yielding 26, 19, and 15 experiments, respectively. For each subset, we evaluated the correlation between the predicted relative tensions of ablated edges and their recoil velocities for both the ForSys baseline and the five StressNET models (evaluated in single-prediction mode).

For StressNET, correlation strength increased monotonically with stricter conditioning thresholds (Supplementary Figure 5). The subset with condition numbers below 500 was used for the final correlation analysis presented in the Results section, as these 15 experiments provided the most stable recoil-velocity estimates.

### 3. Dataset preparation and fine-tuning using Myosin II intensity

#### Dataset and imaging acquisition

To validate StressNET on biological samples, we utilised the dataset used in the ForSys method ^19^, which was obtained by Ventura et al ^28^and consists of 126 microscopy images of the mucociliary epithelium of Xenopus embryos labelled with a non-muscle myosin II A-specific intrabody.

The dataset contained in Ventura et al. ^28^was re-analysed in this context. Stage 16–20 Xenopus embryos expressing the SF9-3xGFP intrabody were imaged using a 3i spinning-disk confocal microscope equipped with a Plan-Apochromat 63×/1.4 NA oil objective, a CSU-X1 spinning-disk head, and an iXon Ultra 888 EM-CCD camera. Maximum-intensity Z projections were generated and segmented with EPySeg. Cell membranes were manually corrected, and membrane outlines were smoothed using a third-order Savitzky–Golay filter (window: 5 pixels). For each membrane, intensities were computed as the mean of vertex-wise values smoothed by a first-neighbour median. To allow comparison with inferred tensions, membrane intensities within each embryo were normalised to a mean of one.

#### Fine-tuning and evaluation using myosin II intensity as ground truth

We used the normalised intensity measurements from the myosin II sensor as the ground truth, as its fluorescence levels at cell-cell interfaces are expected to correlate with membrane stresses. We followed the same preprocessing steps outlined in the ForSys paper and constructed graphs from hand-corrected skeletonised images (Supplementary Figure 3A).

For each image, we computed Pearson correlations between predicted relative stresses and normalised myosin II intensities. For StressNET, correlations were calculated for five independently trained models and averaged to obtain a single performance estimate per sample. Statistical significance was assessed using a paired Wilcoxon signed-rank test.

Each of the five StressNET weight sets was fine-tuned ^31,32^using normalised myosin intensity as the ground truth on randomly selected subsets of 11, 22, 44, and 88 samples from a 70% split of the dataset, with the remaining 30% (38 samples) held out as a static evaluation subset.

To demonstrate that StressNET can be readily fine-tuned out-of-the-box by non-expert users on new datasets with minimal effort, all 20 fine-tuning experiments (5 models × 4 training set sizes) were conducted with preset hyperparameters and a fixed number of training epochs, without relying on early stopping or dynamic learning-rate scheduling.

To evaluate how performance scaled with the amount of fine-tuning data, outliers within each sample-size group were removed using the 1.5×IQR rule, and mean Pearson correlation values were computed for the remaining data. These means were then regressed against the natural logarithm of the number of fine-tuning samples to quantify the trend. The fits again showed agreement with a logarithmic dependence, with coefficients of determination exceeding 0.97 for both single-prediction and augmented-inference variants.

### 4. Extraction and quantitative analysis of tissue-level embeddings

#### Extraction of tissue-level embeddings

We used the “Global Attention Sum Pooling” output from StressNET (Supplementary Figure 1C) as a global embedding for each system. The embeddings were extracted using augmented prediction mode: each system underwent five forward passes with a random in-plane rotation applied to cell coordinates, and the resulting vectors (Global Attention Sum Pooling layer outputs) were averaged to yield a single representation per system. The dataset comprised 100 newly generated simulations per pattern family from the 14 families included during training and the 4 held-out synthetic families (Supplementary Figure 3A), along with samples from in vivo microscopy data (126 Xenopus embryos and 37 zebrafish neuromasts). For robustness, we evaluated the same five sets of model weights trained with different random seeds that were used for the other reported results.

#### Quantitative analysis with PCA and linear probes

All embeddings were standardised using scikit-learn’s StandardScaler (zero mean, unit variance) and analysed separately for each seed. For PCA, we used standardised embeddings to explore the variance structure, and assessed associations between principal component scores and system properties using Spearman’s rank correlation with graph size and point-biserial correlation with the binary “local”/”non-local” label. The PCA model and scaler were fitted exclusively on embeddings corresponding to pattern families included during training, and then applied to all data, including synthetic pattern families excluded from training and in vivo samples. We found that PC1 explained 26.3 ± 1.8 % of the variance of the training data embeddings and PC2 20.6 ± 0.8 %.

To quantify the extent to which global structural information is linearly decodable from the embeddings, we trained linear probes on embeddings corresponding to pattern families included during training using the scikit-learn library with stratified five-fold cross-validation, performed independently for each seed. As a control, we repeated the procedure with 100 random permutations of the labels per fold to estimate the performance expected by chance ^55^. For pattern family prediction, we fit multinomial logistic regression models with L2 regularisation and reported held-out accuracy and confusion matrices.

For graph size prediction, we trained ordinary least squares regressors on the logarithm of the number of nodes and reported R^2^ on the original scale, obtained by back-transforming predictions via exponentiation. As a control, we generated null distributions by repeating each cross-validation experiment on 100 random label permutations per seed.

#### Manifold visualization with UMAP

To further explore the geometric structure of the learned embeddings, we applied UMAP (Uniform Manifold Approximation and Projection) to the Global Attention Sum Pooling vectors across all available data, including synthetic pattern families used during training, held-out synthetic families, and in vivo samples. Embeddings were standardized using scikit-learn’s StandardScaler. UMAP was applied with two output components, Euclidean distance metric, and n_neighbors = 100, while all remaining parameters were kept at their library defaults.

To assess the robustness of the resulting projections to this parameter choice, we evaluated a range of neighborhood sizes (n_neighbors ⍰ {10, 50, 100, 200}). For each configuration, we quantified local class structure in the two-dimensional UMAP space using k-nearest neighbor (k-NN) label agreement, defined as the fraction of a point’s k nearest neighbors in the 2D projection that share its label. Across all tested configurations, k-NN agreement remained stable, ranging from 0.56 to 0.61 for k = 5, 0.55 to 0.60 for k = 10, and 0.53 to 0.58 for k = 20. The magnitude of this variation is limited (absolute changes < 0.06), indicating that local class consistency is largely preserved across a broad range of parameter settings. Importantly, these values are substantially higher than a random baseline (expected score of approximately 0.05 for 20 classes), confirming that the observed neighborhood structure reflects meaningful organization. Qualitative inspection further confirmed that cluster structures and relative relationships between classes were consistent across all tested configurations.

For visualization purposes, we selected n_neighbors = 100 as a representative value balancing local and global structure. In this representation, each point in the two-dimensional projection corresponds to a tissue configuration, including both simulated and experimental samples, colored by pattern family and using marker shape to indicate data source, with circles for training families, crosses for held-out families, and triangles for in vivo samples. The resulting projections enable qualitative visualization of clustering patterns and relationships between embeddings and global system properties across both domains (Figure 4C), complementing the quantitative analyses performed with PCA and linear probes.

## Ethics Statement

Fish used were maintained under standardized conditions. Experiments were performed in accordance with protocols approved by the Ethical Committee of Animal Experimentation of the Helmholtz Zentrum München, the German Animal Welfare act Tierschutzgesetz §11, Abs. 1, Nr. 1, Haltungserlaubnis according to the European Union animal welfare, and under protocol number Gz.:55.2-1-54-2532-202-2014 and Gz.:55.2-2532.Vet_02-17-187 from the “Regierung von Oberbayern” (Germany).

Wild-type *X. laevis* were obtained from Nasco, Wisconsin, Fort Atkinson, WI, USA. The Danish National Animal Ethics Committee has reviewed and approved all animal procedures, housing, and husbandry conditions under permit number 2017-15-0201-01237.

## Data and code availability

StressNET codebase is available on GitHub https://github.com/nicolasaldecoa/StressNET and in Zenodo ^56^.

ForSys is available on GitHub https://github.com/borgesaugusto/forsys and in Zenodo ^57^

## Author contributions

N.A. developed the StressNET code, generated in silico data, inferred *in silico* and in vivo experiments, performed the statistical analysis, quantified the latent space and wrote the article. AB generated in silico data, acquired zebrafish microscopies, analysed in vivo experiments, contributed to statistical analysis, and wrote and edited the article. J.M.-R. acquired the zebrafish microscopy images and analysed the corresponding results. G.V. generated the Xenopus images and edited the article. J.S. supervised the Xenopus project and edited the article. H.L.S. supervised the Zebrafish project and edited the article. O.C. conceived and supervised the project, supervised N.A., and wrote and edited the article.

## Acknowledgments

A.B. acknowledges support through a Research Project Grant from the RPG-2025-081 Leverhulme Trust (awarded to Prof. Ewa Paluch). J.S. acknowledges the support of the Novo Nordisk Foundation (grant number NNF22OC0076414, NNF19OC0056962) and LEO Foundation (grant number LF-OC-19-000219), the European Research Council Consolidator Grant (ERC CoG 101125803 MechanoFate). The Novo Nordisk Foundation Center for Stem Cell Medicine (reNEW) is supported by a Novo Nordisk Foundation grant number NNF21CC0073729. O.C. was funded by a BBSRC grant BB/X014908/1 as well as UADE grants A23T01 and P26T02 to O.C. J.M.-R. is supported by Dirección General de Asuntos del Personal Académico (DGAPA-UNAM) (IA206126). The authors would like to thank Marcos Wappner-Boccaccio, Jack Yu and Anna Foix-Romero for their insightful comments on the manuscript.

