## Supplementary Material for "StressNET: an adaptable deep-learning model for mechanical stress inference in tissues"

[1] Instituto de Tecnología, Universidad Argentina de la Empresa, Buenos Aires, Argentina
[2] Department of Physiology, Development and Neuroscience, University of Cambridge, Cambridge, CB2, UK
[3] Instituto de Neurobiología, Universidad Nacional Autónoma de México (UNAM), Boulevard Juriquilla 3001, Juriquilla, México
[4] Research Institute for Molecular Pathology (IMP), Vienna, Austria
[5] The Novo Nordisk Foundation Center for Stem Cell Medicine (reNEW), University of Copenhagen, Blegdamsvej 3B, 2200 Copenhagen, Denmark
[6] Department of Biomedical Sciences, University of Copenhagen, Copenhagen, Denmark
[7] Division of Science, New York University Abu Dhabi, Saadiyat Island, United Arab Emirates
[8] School of Biosciences, University of Nottingham, Sutton Bonington Campus, Nottingham, LE12, UK
[9] Instituto de Tecnología, Universidad Argentina de la Empresa, Buenos Aires, C1073AAO, Argentina
[&] These authors contributed equally
* Corresponding author: Osvaldo Chara

### Supplementary Results

##### 1. StressNET inference workflow

StressNET is an automated neural network–based pipeline that accurately and precisely infers relative intercellular mechanical stresses from tissue microscopy images (Figure 1). The workflow comprises three stages: (i) extracting tissue geometry as a polygonal 2D mesh, (ii) encoding this geometry as a graph, and (iii) predicting stress distributions via a graph neural network, enabling fast and scalable analysis of complex tissues.

The first stage of the pipeline begins with a microscopy image of an epithelial tissue (Figure 1A-B), which is segmented and skeletonised. The ForSys software is used to convert the 2D segmentation into a polygonal mesh [1] (Figure 1C). Each cell is modelled as a polygon, with edges connecting ordered vertices that delineate the cell boundaries.

The resulting representation serves as the foundation for a graph-based encoding, where nodes correspond to cellular membranes and graph edges connect those nodes that share a junction point (Figure 1D). We frame the problem as a node regression task. Machine learning algorithms such as Graph Neural Networks (GNNs) are well-suited for this task, as they learn node representations via iterative aggregation of information from neighbouring nodes [2]. The underlying rationale is that tension at an intercellular interface is primarily influenced by neighbouring graph nodes rather than by those that correspond to distant cells in the lattice. Therefore, graph convolution operators are an efficient and scalable choice for learning the representations.

We encode the geometrical properties of each cell system into three separate tensors: An adjacency matrix, a Node Features matrix, and an Edge Feature Matrix (Figure 1E-G). The graph encoded as tensors (Figure 1J) serves as input to the StressNET model, which performs node-level regression to predict relative intercellular mechanical stresses (Figure 1K).

#### 2. StressNET models recapitulate state-of-the-art inference results in simulated tissues

To evaluate StressNET's performance relative to existing approaches, we applied it to the benchmark dataset introduced by Borges et al. [1], which consists of vertex-model simulations of epithelial tissues with known ground-truth relative stresses. Although generated independently, the simulations were produced using a procedure similar to that used for our training data. The dataset contains four distinct pattern families, each with 25 samples. Performance was compared against the static ForSys baseline, alongside previously reported DLITE results for the same benchmark.

Relative stresses were predicted using two inference modes: a standard single-prediction mode, consisting of a single forward pass through the network, and an augmented prediction mode. In the augmented mode, nine additional forward passes are performed after applying random rotations to the geometric features, yielding ten predictions in total. These predictions are then averaged and normalised so that their mean equals one to make it comparable with other methods. We quantified performance using a modified score based on MAPE, Pearson's r, and $R^2$ (Materials and Methods). For each sample and modality, we reported performance as the average score across the five models.

We found that across pattern families, StressNET achieved substantially higher scores than the static ForSys baseline in the Circular Furrow and Random Tensions datasets (paired t-tests: $p < 1\times10^{-8}$ and $p < 1\times10^{-10}$, respectively), while no significant differences were observed for the X-axis Furrow (two-sided $p = 0.055$) or Y-axis Furrow (two-sided $p = 0.353$) families. Under augmented inference, StressNET outperformed ForSys in Circular Furrow ($p < 1\times10^{-9}$), Random Tensions ($p < 1\times10^{-10}$), and X-axis Furrow ($p < 1\times10^{-3}$), with no difference for Y-axis Furrow ($p = 0.425$). Augmented inference also significantly improved StressNET's performance over single-pass predictions for all four pattern families (all $p < 1\times10^{-6}$).

These results indicate that StressNET can accurately infer *in silico*-generated data. Moreover, the use of the augmented modality shows that allocating additional computational resources at inference time enhances prediction accuracy (Supplementary Figure 2B; Supplementary Table 1).

#### 3. Assessing StressNET's robustness to segmentation noise

We evaluated StressNET's robustness to input perturbations by introducing Gaussian noise into the in-silico benchmark across four settings, defined by the combination of two noise addition modalities ("intermediate" and "all") and two intensity levels (0.1 and 0.2). Noise intensity was defined relative to local geometry: for each point, the standard deviation of the Gaussian noise was set as a fraction (0.1 or 0.2) of the Euclidean distance to its nearest connected vertex, and noise was sampled accordingly and added to the point's coordinates. In the "intermediate" modality, only the coordinates of non-junction vertices were perturbed, while in the "all" modality, all vertices were affected (Supplementary Figure 2C). For comparison, we used ForSys as a representative systems-of-equations method.

We quantified the degradation in model performance by measuring the relative reduction in our score metric for each sample after introducing noise (Supplementary Figure 2D, Materials and Methods). For each sample, the mean performance across the five StressNET models was reported (Supplementary Table 1). Our results indicate that, on average, StressNET demonstrates significantly greater robustness to geometrical perturbations as compared to the static modality of the ForSys software. When aggregating measurements across all 100 samples, the median relative degradation in our summary score metric is consistently lower for StressNET in every condition ($p < 1\times10^{-14}$ for all conditions; Supplementary Table 2).

These results suggest that StressNET's robustness to geometric noise may offer an advantage for experimental datasets, where imaging limitations are unavoidable, and cell membranes might exhibit irregular geometries that differ from the simplified, regular structures used in computationally generated tissues.

#### 4. Assessing StressNET's performance out-of-distribution

StressNET demonstrated accuracy and precision in its estimations, as well as robustness to noise across the simulation scenarios presented above. However, it is critical that the network generalises well across the diverse topologies encountered in real-world biological applications. Therefore, we evaluated StressNET's performance in terms of mean absolute percentage error (MAPE), using single-prediction and augmented-inference modalities on both in-distribution data (14 simulation families included in training, 100 samples each; 1400 total) and out-of-distribution (OOD) data (four simulation families not included in training, 100 samples each; Supplementary Figure 3A).

We observed a median MAPE for the in-distribution pool was 1.96% for single prediction and 1.63% for augmented inference, while for the out-of-distribution families it was higher, with single-prediction medians of 7.04%, 3.71%, and 2.99%, and augmented-inference medians of 6.32%, 3.33%, and 2.51% for "Radial 1D/2D Inverted", "Trapezoidal X/Y 3-Cells2, and "Trapezoidal X/Y 3-Cells (Small)", respectively (Supplementary Figure 3B).

One-sided Mann–Whitney U tests showed that all OOD families exhibited significantly higher MAPE than the in-distribution pool, with the exception of Furrow Normal (Small), which did not differ significantly (single-prediction: $p = 0.851$; augmented: $p = 0.384$). This result is consistent with the fact that Furrow Normal (Small) shares the same underlying distribution as the in-distribution Furrow Normal simulations, differing only in system size. In contrast, the other three OOD families show significantly greater single-prediction errors ("Radial 1D/2D Inverted": $p = 8.21\times10^{-48}$; Trapezoidal X/Y 3-Cells: $p = 2.01\times10^{-27}$; Trapezoidal X/Y 3-Cells (Small): $p = 6.90\times10^{-11}$). The corresponding augmented-inference comparisons show the same pattern of significance ("Radial 1D/2D Inverted": $p = 4.53\times10^{-47}$; Trapezoidal X/Y 3-Cells: $p = 3.37\times10^{-27}$; Trapezoidal X/Y 3-Cells (Small): $p = 1.09\times10^{-9}$).

Despite higher errors in out-of-distribution families, the overall magnitude remains low (maximum median MAPE of 7.04%), supporting generalisation to previously unseen *in silico* topologies.

##### 5. StressNET benefits from fine-tuning, increasing accuracy on unseen geometries

Deep learning approaches allow freezing a subset of model parameters, and training the remaining ones on a smaller dataset, a process known as fine-tuning [3–6]. Before, we showed that StressNET has a larger error on out-of-distribution data than on in-distribution data. Therefore, we wanted to assess the potential improvements generated through fine-tuning. For this, we selected the out-of-distribution family exhibiting the poorest performance ("Radial 1D/2D Inverted") (Supplementary Figure 4A).

The resulting fine-tuned models showed significant improvements, as measured by the composite score metric (Materials and Methods, Supplementary Table 3). Fine-tuning was performed using datasets of 60, 150, 300, or 600 samples, each split into 75% training and 25% validation data. Performance increased progressively with the amount of fine-tuning data, with the median score in augmented mode rising from 104.4 for the pretrained model to 224.6 after fine-tuning on 600 samples (Supplementary Figure 4B,C). The relationship between performance and the number of fine-tuning samples followed a logarithmic trend, indicating diminishing returns as additional training data were incorporated, consistent with the sublinear convergence behaviour commonly observed in stochastic gradient descent-based optimisation [7].

These results indicate that fine-tuning offers a promising strategy for adapting the model to conditions for which it was not originally trained. The consistent gains observed in out-of-distribution simulations suggest that this approach could be effective when StressNET is applied to real experimental data, where pronounced domain shifts are expected.

### Supplementary Materials and Methods

##### Synthetic Dataset Generation

We developed a suite of Python tools that interface with the SurfaceEvolver [8] software through the SeapiPy package [1,9]. Using these tools, we generated 18 distinct scripts, each corresponding to different types or "families" of simulations. In each case, the scripts sample from a predefined parameter space to establish the system's initial conditions, such as the number of cells, their shapes, and other relevant geometric properties.

A Voronoi tessellation is used to generate a lattice, after which tensions are assigned to the edges based on patterns characteristic of each simulation type. The resulting configurations are converted into files containing definitions and instructions compatible with SurfaceEvolver, which is then used to simulate the tissue deformation under the forces applied along the edges. Simulations run for a sufficient number of steps to allow the system to reach an equilibrium state (Supplementary Figure 2A).

Across the 18 families, simulations differed along multiple design dimensions, including the parameter ranges sampled during initialisation and the patterns used to assign target edge tensions prior to relaxation. Although these variations were not limited to a single factor, the families can be broadly grouped according to how tensions were specified. In a subset of families ("local"), edge tensions were defined as smooth functions of spatial position, resulting in spatially continuous stress distributions. In contrast, in "non-local" families, tensions were sampled independently from predefined probability distributions without reference to spatial coordinates. Families also differed in the range of sampled cell numbers, thereby introducing additional variability in graph size and overall structural scale.

To evaluate the model's generalisation capacity, 4 out of the 18 simulation families were excluded from the training and validation sets and reserved exclusively for the test set. For each of these families, 300 samples were generated, while 2,000 samples were produced for each of the remaining 14 families. This entire process was conducted twice, with the number of vertices defining each cell interface set to either 5 or 9, labelled as "5v" and "9v", respectively.

After conducting experiments to compare performance across tasks between models with different vertex counts and testing methods to adapt inputs for inference, we expanded the 9v dataset by upscaling the 5v dataset's edge features using degree-2 spline interpolation. We split the in-distribution simulations into 75% for training, 10% for validation, and 15% for testing. This resulted in a training set of 21,000 samples for the 5v models and 42,000 for the 9v models, with 2,800 and 5,600 samples used for validation, respectively. After adding the out-of-distribution samples, the test set contained 5,400 samples for 5v models and 10,800 for 9v models, which were used to compare performance for hyperparameter tuning and model selection.

#### Training StressNET on Synthetic Data

Models were implemented in TensorFlow [10] and trained using the Adam optimiser [11] with a learning rate of 3e-4 and gradient clipping at 1.0. The batch size was set to 32, and training used the mean squared error as the loss function.

To improve generalisation, we applied on-the-fly data augmentation [12] to the edge features, including random rotations (always applied) and random jitter (applied with a 25% chance to each sample). The learning rate was adaptively reduced by a factor of 0.5 when the validation loss failed to improve over 5 consecutive epochs, with a minimum rate of 1e-6. Training was stopped early if no improvement was observed on the validation set over more than 10 epochs, and the best-performing weights were restored.

To assess the stability of the training process and improve robustness in subsequent evaluations on synthetic and experimental data, each experiment was repeated with five different random seeds.

#### StressNET models recapitulate state-of-the-art inference results in simulated tissues

To benchmark StressNET against existing approaches, we used the synthetic dataset introduced by Borges et al. [1], which comprises four pattern families—Circular Furrow, Random Tensions, X-axis Furrow, and Y-axis Furrow—each containing 25 samples generated using a vertex model implemented in SurfaceEvolver through SeapiPy [1,9]. The ForSys engine was used to generate the 2D lattice for each sample, which we converted into a graph representation for use with StressNET. Predictions of relative edge tensions were obtained in two modes: a single-pass inference (one forward pass through the network) and an augmented inference mode in which nine additional forward passes were performed after applying random rotations to the geometric features; the ten resulting tension vectors were averaged and renormalised to yield a mean of 1.

For performance assessment, we employed a modified version of the scoring function from Borges et al. [1], which combines Mean Absolute Percentage Error (MAPE), Pearson's correlation coefficient (r), and the coefficient of determination ($R^2$). To prevent any single metric from dominating the score, input values were saturated at 1% for MAPE and 0.99 for r and $R^2$. To ensure robustness to model initialisation, five independently trained StressNET models (identical training settings, differing only in random seed) were evaluated, and the reported score for each sample corresponds to the mean across these five instances. Statistical comparisons were conducted using paired t-tests. For each pairwise contrast, the paired differences were assessed using the Shapiro–Wilk tests prior to paired t-tests. Although some comparisons showed mild deviations from normality, the paired t-test is robust to such deviations for moderate sample sizes. All statistical analyses were performed in Python using SciPy (v1.13.1).

#### StressNET benefits from fine-tuning, increasing accuracy on unseen geometries

During fine-tuning, approximately 97% of StressNET's parameters were frozen [4], leaving only the weights of the final two fully connected layers and the intermediate LayerNorm rescaling vector trainable (Figure 3C; Supplementary Figure 1D). Fine-tuning was conducted on subsets of 60, 150, 300, and 600 samples from a newly generated set of the out-of-distribution simulation family "Radial 1D/2D Inverted", leaving the original set of 100 samples for evaluation. Within each subset, 25% of the data was reserved for validation. Models were optimised using the Adam optimiser with a learning rate of 0.001 and gradient clipping at 5.0, minimising the mean squared error. During training, on-the-fly data augmentation was applied to the edge features in the form of random rotations. The learning rate was reduced by a factor of 0.5 if the validation loss did not improve for 3 consecutive epochs, down to a minimum of $1\times10^{-6}$. Training was stopped early if no improvement was observed on the validation subset over more than 6 epochs, and the best-performing weights were restored.

To evaluate how performance scaled with the amount of fine-tuning data, we quantified the relationship between the mean score and the number of fine-tuning samples (60, 150, 300, and 600). For each sample-size condition and for each inference mode, outliers were removed using the 1.5×IQR criterion applied independently within each group of replicates. The remaining score values were averaged per sample size, and the natural logarithm of the corresponding sample count was used as the predictor in a linear regression model. The resulting fits captured nearly all of the variance in the data ($R^2 > 0.979$ across modes), indicating that performance followed an approximately logarithmic scaling trend with increasing amounts of fine-tuning data.

### Supplementary Figures

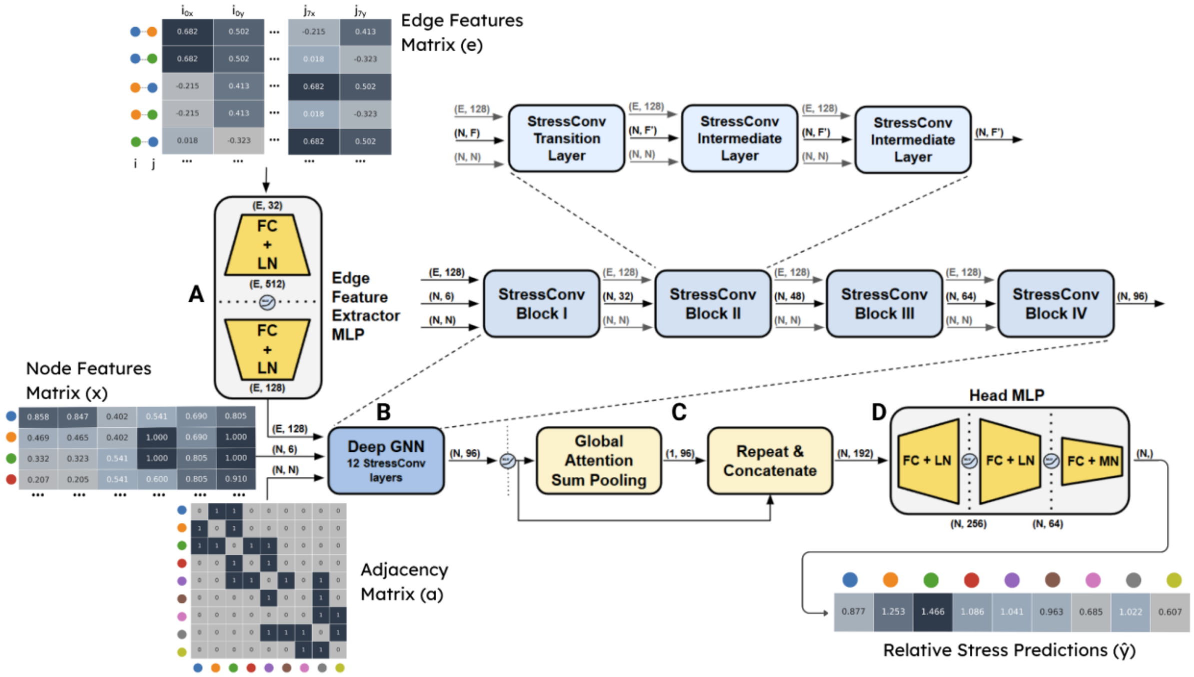


**Supplementary Figure 1. StressNET architecture for learning relative intercellular tension from tissue graphs.** StressNET consists of four main components, each fulfilling a distinct role within an end-to-end trainable architecture. **(A)** An edge feature extractor MLP embeds raw geometric edge features into a higher-dimensional representation. **(B)** A deep graph neural network composed of multiple StressConv layers integrates node and edge information, progressively updating node embeddings. **(C)** A global attention-based pooling layer aggregates all node representations into a single tissue-level descriptor, which is then concatenated back to each individual node embedding. **(D)** A final MLP head performs node-level regression to predict the relative magnitude of intercellular tension at each interface.

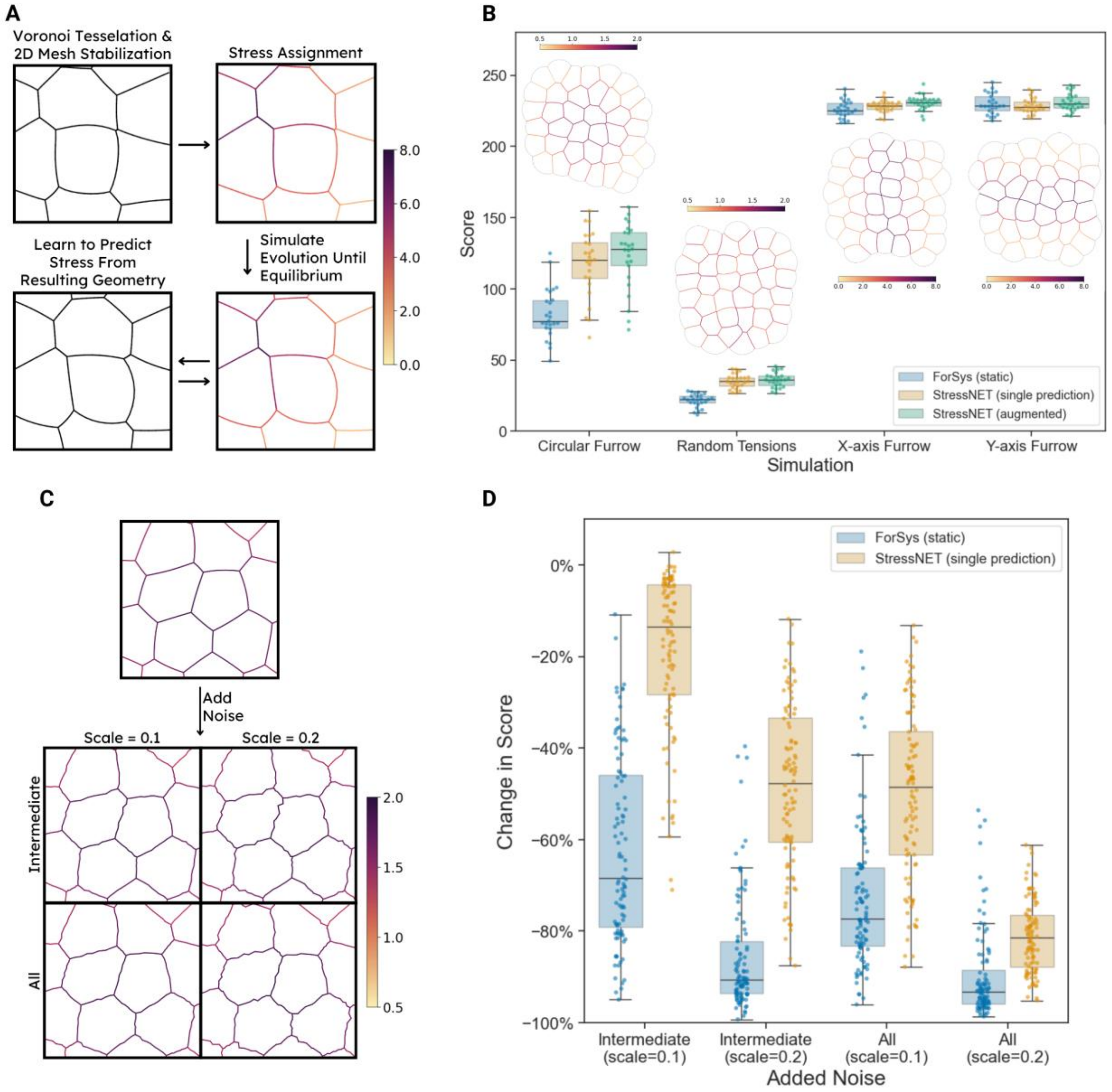


**Supplementary Figure 2. StressNET outperforms existing methods and generalises across synthetic and biological tissues.** **(A)** Synthetic benchmark generation follows the ForSys static pipeline: a 2D lattice is created via Voronoi tessellation, interfacial tensions are assigned, and tissue geometry is stabilised and evolved to mechanical equilibrium using SurfaceEvolver. The resulting lattices are used to train StressNET base models. **(B)** Performance comparison of ForSys (static), StressNET (single prediction), and StressNET (augmented) across four benchmark simulation families. StressNET consistently outperforms ForSys in the Circular and Random Tensions families, and its augmented variant further improves performance. Insets show representative samples from each simulation family. **(C)** Schematic of the perturbation protocol used to evaluate robustness. Gaussian noise is added to vertex coordinates at two intensity levels (0.1 and 0.2), either to non-junction vertices only (“intermediate”) or to all vertices. **(D)** Performance degradation under geometric noise. StressNET maintains significantly higher accuracy than ForSys across all noise conditions, demonstrating greater robustness.

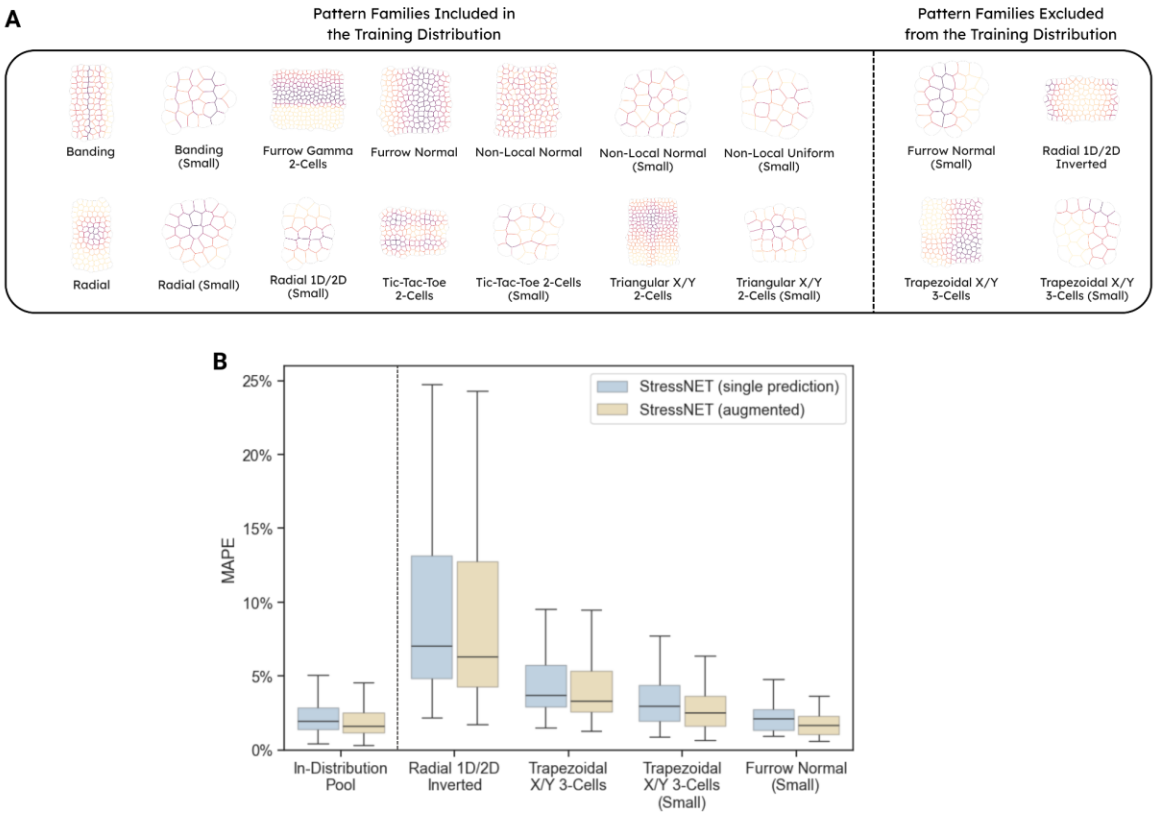


**Supplementary Figure 3. StressNET performance on in- and out-of-distribution simulation families. (A)** Representative examples of simulation families included in the training set (left) and families excluded from training (right). **(B)** Boxplots of the mean absolute percentage error (MAPE) for StressNET predictions on in-distribution and out-of-distribution data, shown for single-prediction (blue) and augmented-inference (yellow) modalities. Out-of-distribution errors are higher than in-distribution ones, as expected, with median MAPE rising from roughly 1.6-2.0% in-distribution to 3-7% across unseen pattern families. For all boxplots, whiskers extend to the most extreme data points within 1.5× the interquartile range.

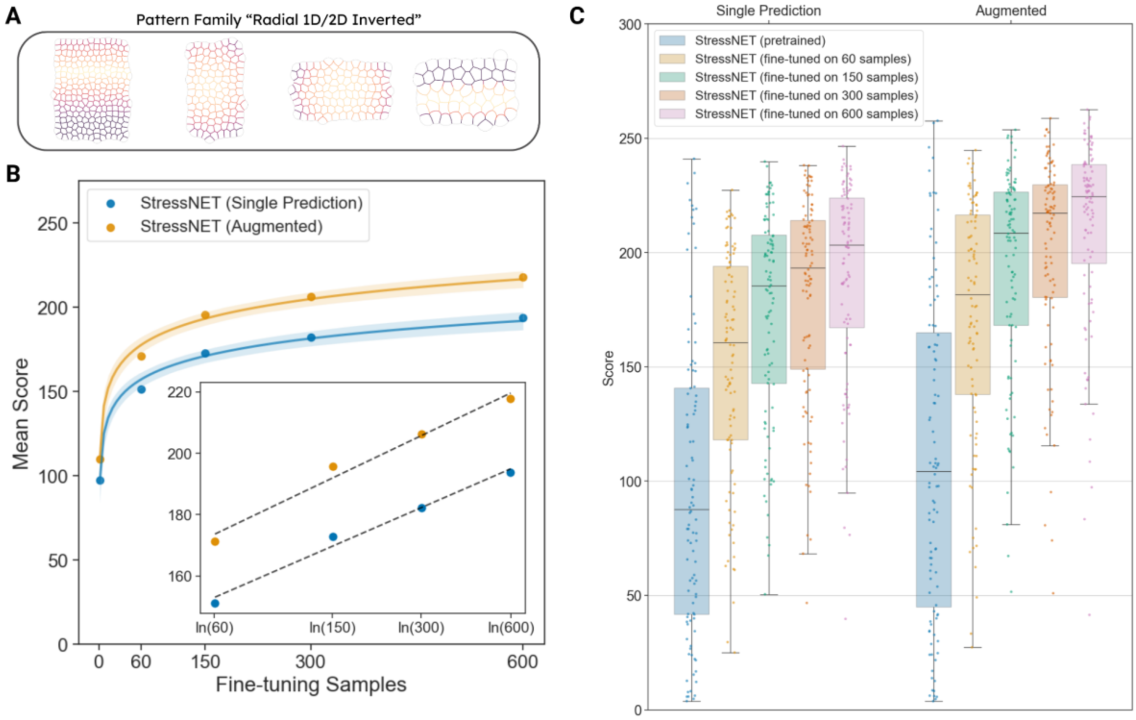


**Supplementary Figure 4. StressNET fine-tuning on out-of-distribution synthetic data. (A)** Visualised samples from the out-of-distribution family 'Radial 1D/2D Inverted,' which exhibited the highest median MAPE (7.04%) among all out-of-distribution families. **(B)** Mean scores as a function of the number of fine-tuning samples follow a logarithmic trend. The shaded regions represent confidence intervals derived from bootstrap resampling. The inset presents the linear regression on the mean scores computed in log-transformed sample space. **(C)** Distribution of scores for different fine-tuning sample sizes and inference modalities, illustrating that average performance improves monotonically with additional fine-tuning data. Boxplot's whiskers extend to the most extreme data points within 1.5× the interquartile range.

A

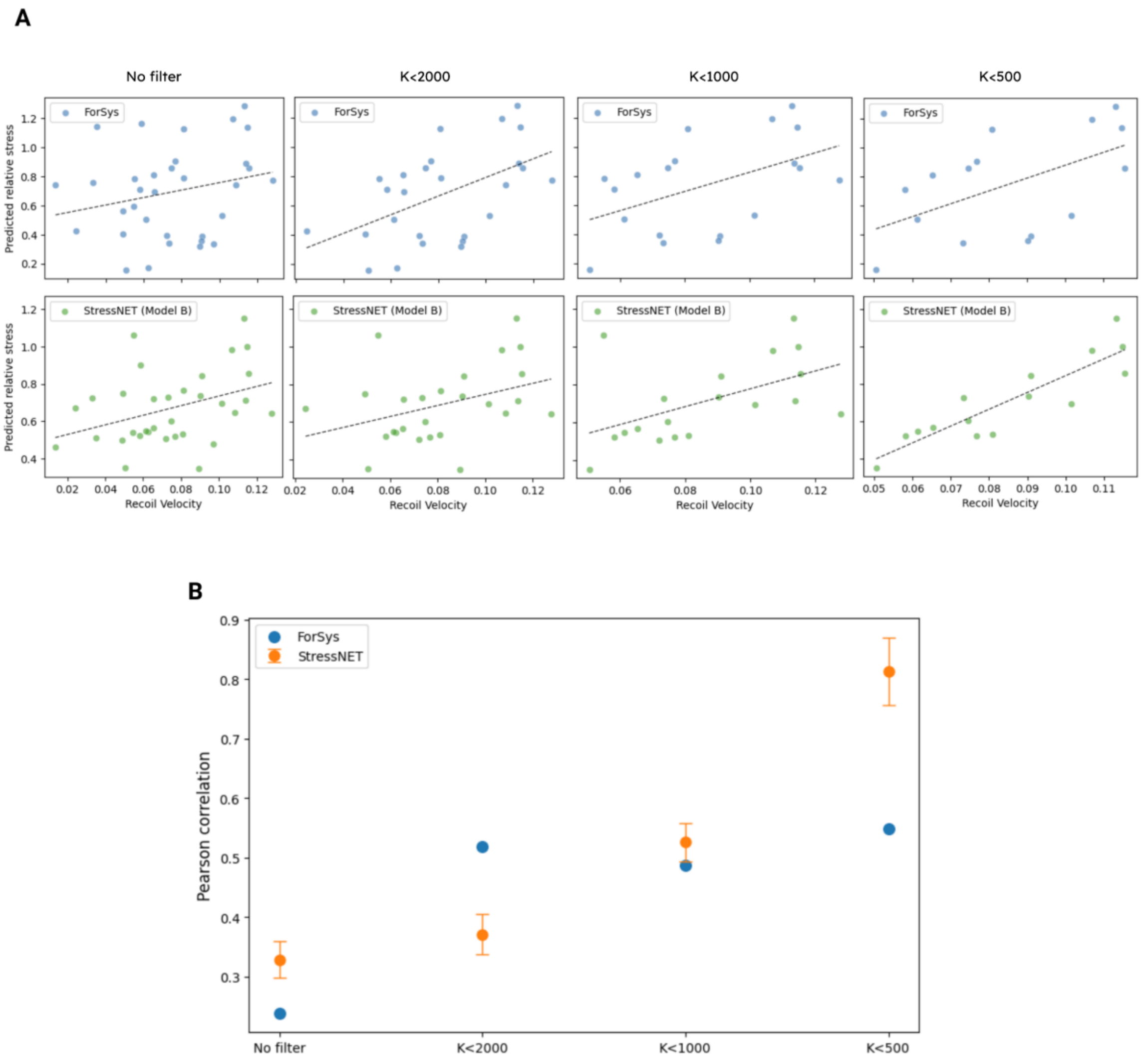


***Supplementary Figure 5**. Correlation between predicted tensions and recoil velocities improves with increasingly stable recoil-velocity estimates.*

*(A) Scatter plots of estimated recoil velocity versus predicted relative tension for ForSys (top row) and one representative StressNET model (bottom row), shown across progressively more stringent subsets defined by the maximum allowed condition number of the linear fit used to estimate recoil velocity (no filter; K < 2000; K < 1000; K < 500). The dotted lines indicate linear fits illustrating the trend within each subset. As increasingly unstable fits are removed, the tension–recoil relationship becomes clearer for both models. (B) Pearson correlation coefficients between predicted relative tensions and recoil velocities computed on the same progressively filtered subsets. Points show the correlation for ForSys and the mean ± s.d. across the five StressNET base models. Correlation strength for StressNET increases monotonically with stricter conditioning thresholds, reflecting improved numerical stability of the recoil-velocity estimates.*

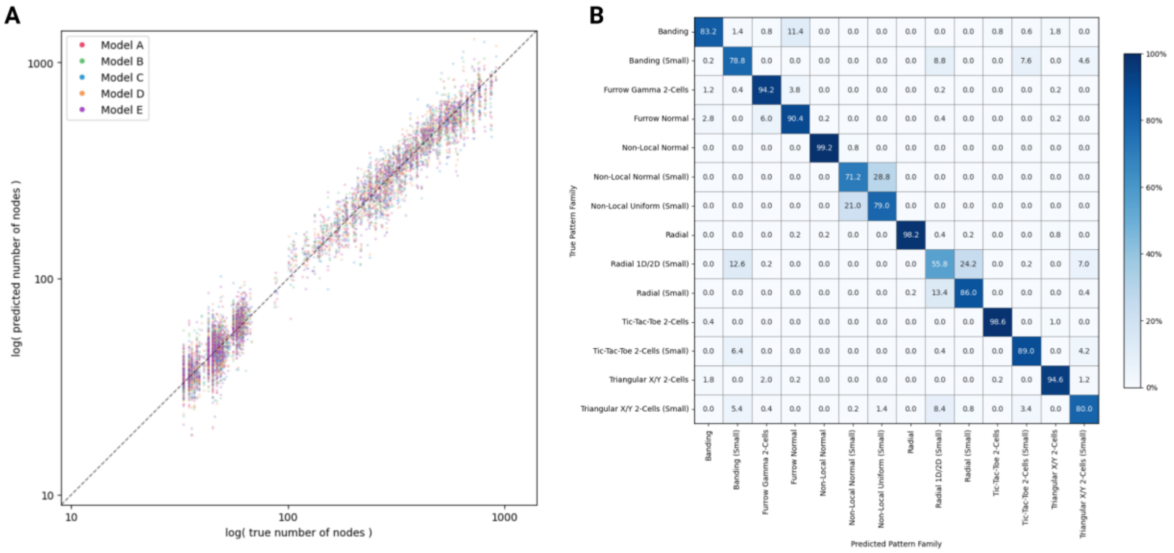


***Supplementary Figure 6***. *Characterisation of StressNET latent representations.*

*(A) Log–log scatter plot comparing predicted versus true numbers of nodes for linear regression probes trained to predict the logarithm of graph size using the same cross-validation setup. Each point represents one sample, colored by model replicate (A-E). The dashed line indicates the identity line (y = x). (B) Averaged confusion matrix of logistic regression probes trained to classify pattern families from the Global Attention Sum Pooling embeddings, using 5-fold cross-validation across five independently trained StressNET models. Values indicate mean out-of-fold prediction accuracy (%) for each true–predicted family pair.*

### Supplementary Tables

| Model | Score (mean ± std) | | | |
|---|---|---|---|---|
| | **Circular Furrow** | **Random Tensions** | **X-axis Furrow** | **Y-axis Furrow** |
| ForSys (static) | 82.4 ± 17.9 | 21.4 ± 4.7 | 226.1 ± 6.1 | 229.8 ± 7.1 |
| StressNET | 117.6 ± 23.0 | 35.0 ± 5.0 | 228.3 ± 4.2 | 228.6 ± 5.1 |
| StressNET (augmented) | **123.6 ± 23.0** | **35.9 ± 5.1** | **230.8 ± 5.1** | 231.0 ± 6.0 |

**Supplementary Table 1.** Performance comparison between StressNET and ForSys across four synthetic datasets. StressNET significantly outperformed static ForSys on Circular Furrow and Random Tensions, with no significant difference on X-axis and Y-axis Furrow. Augmented inference further improved StressNET performance on Circular Furrow, Random Tensions, and X-axis Furrow, while showing no significant change on Y-axis Furrow. Across all datasets, augmented inference significantly outperformed single predictions.

| Model | Relative Regression Score Reduction (median [$Q_1$, $Q_3$]) | | | |
|---|---|---|---|---|
| | **Intermediate (0.1)** | **Intermediate (0.2)** | **All (0.1)** | **All (0.2)** |
| ForSys | 0.68 [0.46, 0.79] | 0.91 [0.82, 0.94] | 0.77 [0.66, 0.83] | 0.93 [0.89, 0.96] |
| StressNET | **0.14 [0.04, 0.28]** | **0.48 [0.33, 0.60]** | **0.48 [0.36, 0.63]** | **0.81 [0.76, 0.88]** |

**Supplementary Table 2.** Median relative regression score reduction (with first and third quartiles) for StressNET and ForSys under four noise conditions. Values represent degradation in the score metric relative to noise-free inputs, with lower values indicating greater robustness. StressNET consistently showed significantly lower degradation than ForSys in all conditions.

| Model | Score (median [$Q_1$, $Q_3$]) | MAPE (%) (median [$Q_1$, $Q_3$]) |
|---|---|---|
| StressNET<br>Pre-trained (single prediction) | 87.7 [41.8, 140.8]<br>(n=100) | 7.0 [4.8, 13.2]<br>(n=100) |
| StressNET<br>Pre-trained (augmented) | 104.4 [45.1, 165.0]<br>(n=100) | 6.4 [4.4, 12.7] (n=100) |
| StressNET<br>fine-tuned using 600 samples<br>(single prediction) | 203.3 [167.3, 223.9]<br>(n=100) | 3.6 [2.8, 4.3]<br>(n=100) |
| **StressNET**<br>**fine-tuned using 600 samples**<br>**(augmented)** | **224.6 [195.3, 238.6]**<br>(n=100) | **2.9 [2.3, 4.0]**<br>(n=100) |

**Supplementary Table 3.**
Performance of StressNET on a representative out-of-distribution simulation family before and after fine-tuning. Scores are reported as the median and interquartile range across 100 evaluation samples, measured using the modified Score from Borges et al. In addition, median MAPE is provided to facilitate comparison with the pre-trained model's performance on the in-distribution evaluation pool (1.96% for single prediction, 1.63% for augmented inference). Fine-tuning substantially improved performance, reducing the median MAPE from 7.04% (pre-trained, single prediction) and 6.37% (pre-trained, augmented) to 3.59% and 2.90%, respectively.
